# ISG15/USP18/STAT2 is a molecular hub regulating autocrine IFN I-mediated control of Dengue and Zika virus replication

**DOI:** 10.1101/784678

**Authors:** Constanza Eleonora Espada, Edroaldo Lummertz da Rocha, Taissa Ricciardi-Jorge, Adara Aurea dos Santos, Zamira Guerra Soares, Greicy Malaquias, Daniel Oliveira Patrício, Edgar Gonzalez Kozlova, Paula Fernandes dos Santos, Juliano Bordignon, Thomas J. Sanford, Teodoro Fajardo, Trevor R. Sweeney, André Báfica, Daniel Santos Mansur

**Affiliations:** Laboratório de Imunobiologia, Departamento de Microbiologia, Imunologia e Parasitologia, Centro de Ciências Biológicas, Universidade Federal de Santa Catarina, Santa Catarina CEP 88040-900, Brazil; Laboratório de Virologia Molecular, Instituto Carlos Chagas, ICC/Fiocruz-PR, Curitiba, Paraná, CEP 81350-010, Brazil; Division of Virology, Department of Pathology, University of Cambridge, Addenbrooke’s Hospital, Hills Road, Cambridge, CB2 0QQ, UK

**Keywords:** Dengue virus, Zika virus, ISG15, USP18, type one interferon, ISGylation, antiviral response, immune evasion, NS5, STAT2

## Abstract

The establishment of a virus infection is the result of the pathogen’s ability to replicate in a hostile environment generated by the host’s immune system. Here, we found that ISG15 restricts Dengue and Zika viruses’ replication through the stabilisation of its binding partner USP18. ISG15 expression was necessary to control DV replication driven by both autocrine and paracrine type one interferon (IFN-I) signalling. Moreover, USP18 competes with NS5-mediated STAT2 degradation, a major mechanism for establishment of flavivirus infection. Strikingly, reconstitution of USP18 in ISG15-deficient cells was sufficient to restore the STAT2’s stability and restrict virus growth, suggesting that the IFNAR-mediated ISG15 activity is also antiviral. Our results add a novel layer of complexity in the virus/host interaction interface and suggest that NS5 has a narrow window of opportunity to degrade STAT2, therefore suppressing host’s IFN-I mediated response and promoting virus replication.

**IMPORTANCE:** Disease is an emergent property that results from a microorganism’s ability to replicate in a given host and the latter’s immune response. Here we describe how the immunoregulatory function of ISG15 (an interferon stimulated gene) affects Dengue and Zika virus replication by occupying a niche used by the virus non-structural protein 5 (NS5) to evade host’s immunity. In the absence of ISG15, NS5 efficiently degrades a main signalling hub of innate immunity (STAT2), leading to cell immune suppression and consequently virus growth. This sheds light into how flaviviruses intimately interact with the host immune system and could lead to a host-based therapy target.

## INTRODUCTION

Cells detect infection by recognizing molecular patterns derived from pathogen’s constituents (PAMPs) or cell damage (DAMPs). During viral infection, nucleic acid is a major signal that triggers the innate immune response, inducing a type one interferon (IFN-I)- mediated antiviral state [1, 2]. IFN-I binds to its cognate receptor and activates the JAK/STAT pathway, leading to expression of hundreds of interferon stimulated genes (ISGs) that make the intracellular environment hostile to viral replication in infected and proximal cells [3]. The evolutionary arms race between viruses and its hosts led to evolution of immune evasion mechanisms that are crucial for successful viral replication. Considering IFN-I’s importance in viral infection control, many immunomodulatory proteins target this signalling pathway [4].

Several flaviviruses, such as dengue virus (DV), Zika virus (ZIKV) and yellow fever virus (YFV), have emerged and re-emerged over recent years and are the leading cause of human arbovirus infection [5, 6]. DV alone infects nearly 400 million people every year [7] with extensive health and economic burden [8]. A requirement for effective flavivirus emergence is the ability to counteract the human immune system.

The compact flavivirus genome encodes seven non-structural proteins that are responsible for viral replication and immune evasion. Six of these proteins are not secreted implying that intracellular pathways are central targets for evasion [9–13]. For instance, DV non-structural protein 5 (NS5), which is the viral RNA-dependent-RNA-polymerase (RdRp) and a methyltransferase, mediates STAT2 degradation by facilitating its interaction with UBR4, leading to its ubiquitination and subsequent proteasomal targeting [14]. This evasion pathway is functional in humans but not mice due to differences in the amino acid sequence of human and murine STAT2 [15].

ISG15 is an intracellular and secreted ubiquitin-like protein that has three described functions. Extracellular ISG15 acts as a cytokine, leading to the expression of IFNγ and IL-10 in diverse immune cells [16–18]. It has been suggested that humans lacking ISG15 have severe mycobacterial disease due to deficiency in IFNγ production by NK cells [19]. Moreover, ISG15 mRNA is highly expressed in active tuberculosis and strongly correlates with disease severity [17, 20].

ISG15 is conjugated to other proteins through a three-step ubiquitination-like process [21] in which the main ligase for ISGylation is the HECT domain and RCC1-like domain-containing protein 5 (HERC5) [22, 23]. ISGylated proteins are affected in several different ways, such as increased or reduced stability and activity. ISG15 can also be conjugated to viral proteins, impacting their function, and therefore belong in the plethora of ISGs with a direct antiviral function [3, 24].

The third and more recently described role of ISG15 is its IFN-I modulatory function. Non-conjugated ISG15 binds and stabilises the ISG USP18, a protease that negatively regulates IFN-I signalling and also serves as a deISGylation protein [25–27]. Correspondingly, individuals lacking ISG15 are prone to severe interferonopathies due to decreased USP18 function and increased IFN-I signalling. Interestingly, ISG15-deficient patients do not have enhanced susceptibility to viruses suggesting ISG15 is not necessary to control ubiquitous viral infections in vivo [28]. In contrast to the indirect role of ISG15 in negative regulation of IFN-I signalling through the USP18 axis in humans [28, 29], murine ISG15 blocks replication of human viruses such as Influenza and HSV-1[30]. These findings implicate ISG15 as an important molecule inhibiting IFN-I-mediated actions but also suggest that ISG15 may mediate host cell intrinsic mechanisms triggered by viruses. However, how ISG15 bridges these apparently two paradoxical phenomena is unclear. Specifically, it is possible that ISG15 directly regulates proteins exploited by viruses during early intracellular infection events.

Here, we observed that, unlike several other ISGs, ISG15 is highly expressed in infected cells containing the DV genome. Furthermore, independently of its ISGylation function, ISG15 restricts flavivirus replication primarily in the infected cell by stabilising USP18, which in turn competes with viral NS5 for binding to STAT2. Our results suggest that flaviviruses exploit an ISG15-mediated IFN-I regulatory mechanism to evade innate immunity and enable replication.

## RESULTS

### ISG15 is expressed in DV-infected cells

The cell is a fundamental unit for viral infection control and developments in single-cell sequencing technology have enabled examination of host-pathogen interactions in great detail. Zanini and colleagues generated single-cell RNA sequencing data from human cells (PBMCs and the HuH7 hepatoma cell line) infected with DV [31, 32]. We re-analyzed these available single-cell transcriptomic data dividing cells into three categories: uninfected, infected and bystander. Here we define uninfected cells as those derived from healthy donors; bystander cells as those derived from an infected patient or have been exposed to the virus but did not have the viral RNA detected and infected cells as those in which viral genome was detected. We used t-Distributed Stochastic Neighbour Embedding (tSNE) analysis to visualise cell-to-cell relationships in space of reduced dimensionality. As reported previously [31, 32], global cellular mRNA expression profiling was not sufficient to separate infected or bystander from uninfected cells, suggesting a high variability of gene expression in these samples (Figure 1A and S1A). As IFN-I are key elements in controlling infection, we filtered the results of the differential gene expression analysis using the gene ontology (GO) term for “type one interferon”. The Venn diagram in figure 1B shows that from the 394 differentially expressed (DE) genes in peripheral blood mononuclear cells (PBMCs) of patients infected with DV or healthy donors, 37 were ISGs (IFN-I GO). ISG15, UBE2L6, HERC5 and USP18, members of the ISGylation pathway, were differentially expressed during DV infection (Figure 1C underlined). Interestingly, this is in contrast to other single cell experiments using Influenza virus as a model, where all members of the ISGylation pathway seem to be expressed at similar levels in both infected and bystander cells [33], HERC5 was the only member of the ISGylation pathway with a higher expression level in bystander cells (Figure 1C and D). In the data set derived from the HuH7 cell line, ISG15 was the only canonical antiviral protein expressed in DV genome-containing cells (Figure S1C and D). This is in agreement with previous reports that the Huh7 cell line does not produce IFN-I upon viral infection [34–36] and could explain the high number of infected cells (Figure S1A) in comparison with the number of infected PMBCs (Figure 1A). In PBMCs, NK, monocytes and B cells were the infected cells with higher ISG15 expression and similar to the data from Zanini and colleagues [32] B cells and monocytes being proportionately the most infected cells (Figure 1E). These results show that in contrast to most ISGs, ISG15 and other components of the ISGylation pathway are enriched in cells where the DV genome was present.

**Figure 1.**
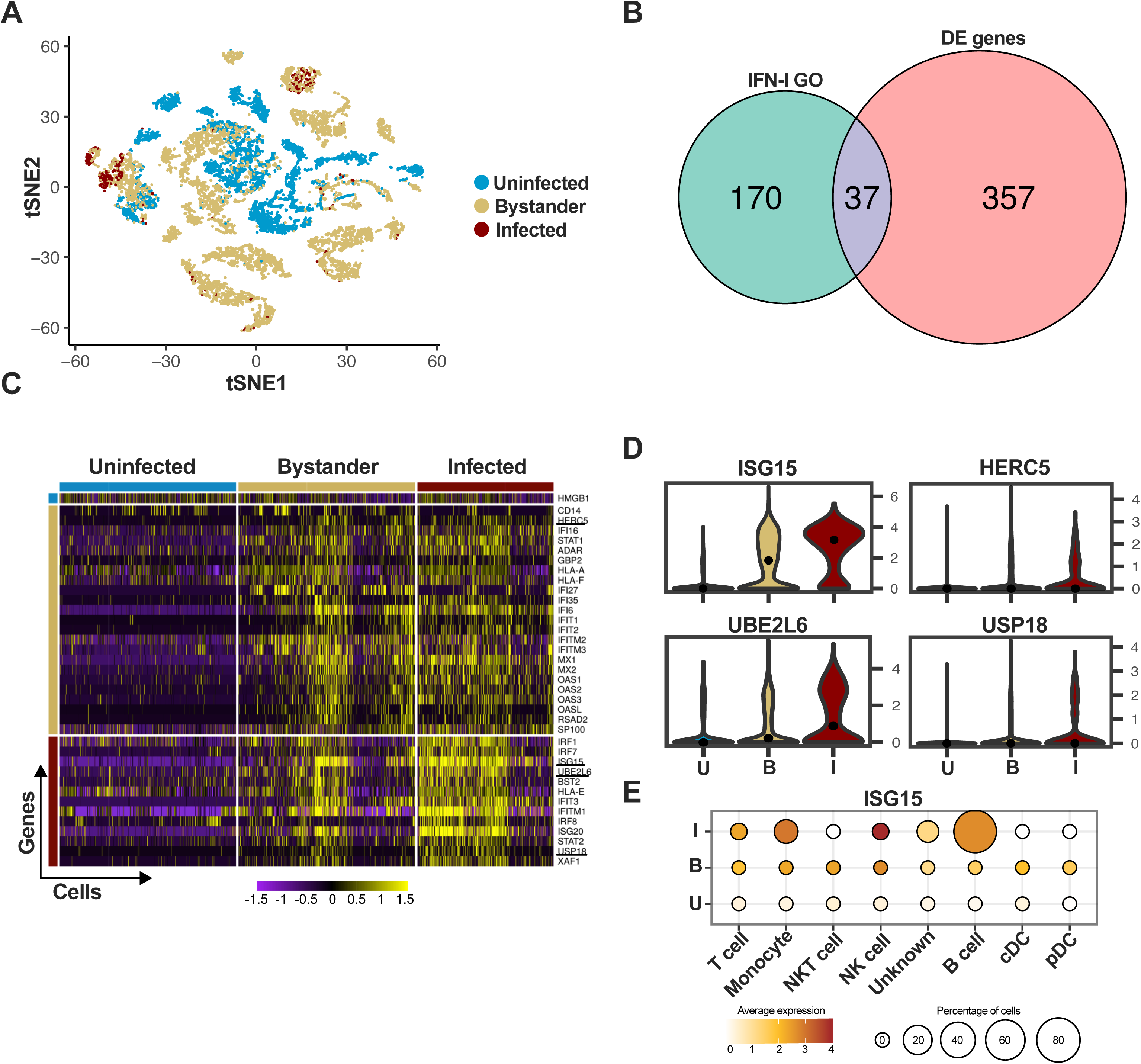
ISG15 is expressed in DV-infected PBMC. (A) tSNE was used to visualise the single-cell global transcriptome data. Blue dots represent uninfected cells, derived from healthy donors. Beige dots represent bystander cells and red dots represent infected cells, both derived from patients infected with DV. (B) Differential expression of ISGs in PBMCs of patients infected with DV. The Gene Ontology term “type one interferon” was used to filter the results from the single-cell RNA sequencing. (C) Single cell ISG expression variability in uninfected, bystander and infected. ISGylation related genes are underlined. Blue bar represents uninfected cells, derived from healthy donors. Beige bar represents bystander cells and red bar represents infected cells, both derived from patients infected with DV. (D) Violin plot representing the expression of ISGylation family members in uninfected [U], bystander [B] and infected [I] PMBC. (E) Expression of ISG15 in DV infected PMBCs. Size is proportional to the percentage of infected cells in each cell type. Colour intensity represents ISG15 average expression.

### ISG15 restricts DV and ZIKV replication

The enrichment of ISG15-related genes at a single-cell level led us to investigate how ISG15 might impact flavivirus replication in a human cell. We used an A549 cell line lacking ISG15, previously generated using CRISPR/Cas9 in our lab [17]. A549 cells were chosen due to their ability to support flavivirus replication and most importantly to produce and respond to IFN-I [37–40].

ISG15-deficient cells were more susceptible to DV infection as shown by an increased foci size (Figure 2A and B) and number of infected cells per foci (Figure 2C). In addition, we determined the kinetics of DV replication and dissemination by using a low multiplicity of infection (MOI), to allow for viral spread through secondary infection events. Percentages of infected cells over time (Figure 2D), relative DV Pre-Membrane RNA quantification in the supernatant (Figure 2E) and infectious particle formation (Figure 2F) were also increased in the absence of ISG15. DV infection at 4 °C for 2 hours followed by relative intracellular DV genome quantification indicated no differences in viral RNA between WT and knockout cells up to 48 hours post-infection (Figure 2G), indicating that ISG15 plays a role in the DV life cycle at a stage after viral entry. Importantly, reconstitution of ISG15 expression in knockout cells led to phenotypic reversion (Figure 2H and I), confirming that the effects observed in our experiments were caused by depletion of ISG15.

**Figure 2.**
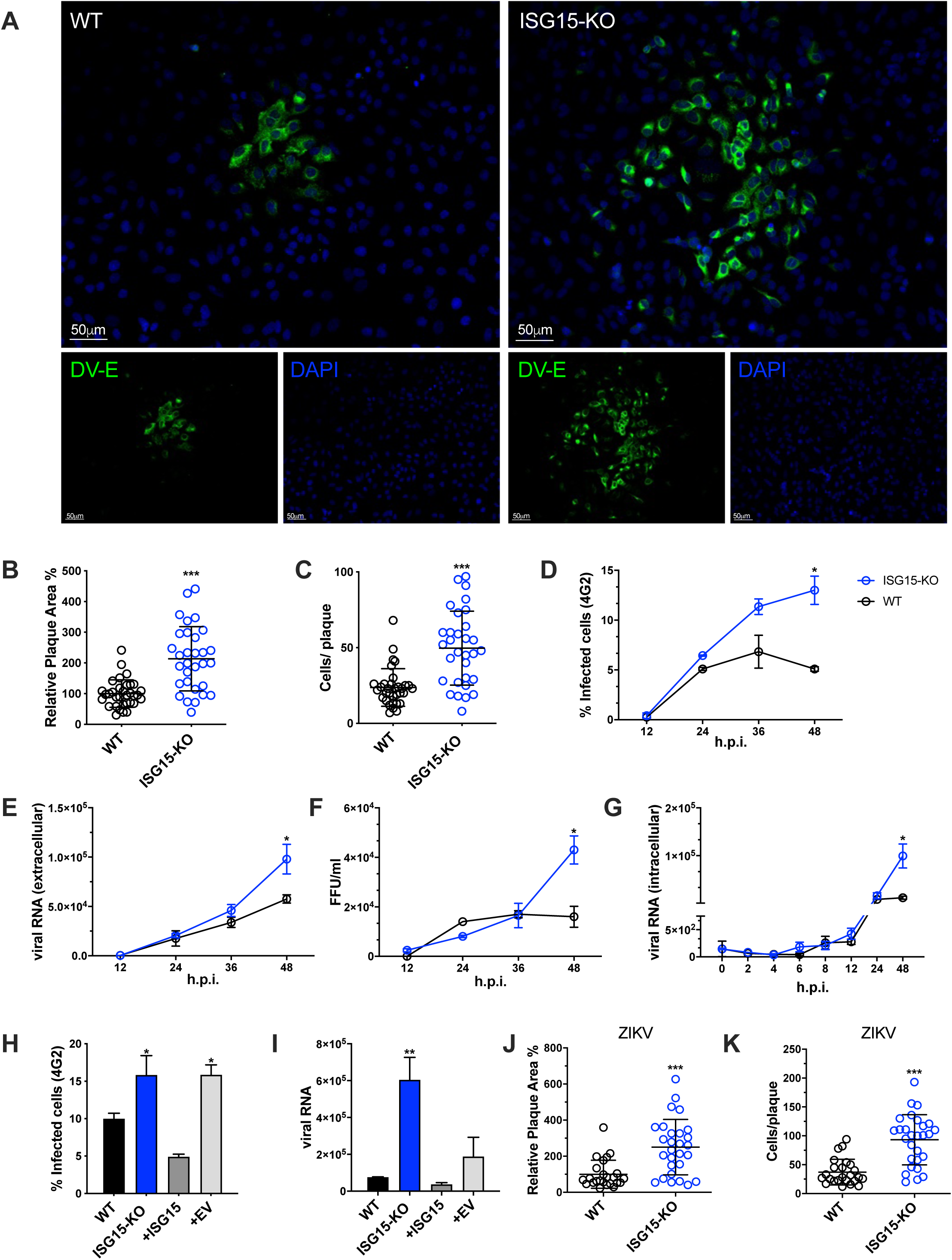
ISG15 restricts DV and ZIKV replication. (A) A549 WT and ISG15 KO were infected with 20 DV PFUs. At 36 hpi cells were fixed, permeabilized and stained for the flavivirus E protein using 4G2 antibody. Displayed images were acquired with a Leica DMI6000 B microscope. (B and C) DV relative foci area (B) and the number of infected cells per foci (C) quantified by ImageJ software and analysed using Welch’s t test. Error bars represent mean ± SD. Results are representative of three independent experiments. (D, E and F) Multiple-step DV growth curve in A549 cells. Cells were infected at an MOI of 0.01 and harvested at multiple time points. Shown is the percentage of cells infected as measured by E protein staining (4G2+) (D), extracellular viral RNA relative expression by RT-qPCR (E) and titration by focus forming assay (FFA) (F). Statistical analyses were conducted using unpaired t tests. Error bars represent mean ± SD. Results are representative of three independent experiments. (G) Changes in viral RNA relative expression over time following binding of DV to A549 WT and ISG15 KO cells at 4°C and analysed using unpaired t test. Error bars represent mean ± SD. Results are representative of two independent experiments. (H and I) Complementation of A549 ISG15 KO cells with ectopically expressed ISG15. Cells were infected at an MOI of 0.01 and at 36 hpi cells were fixed and processed for measurement by flow cytometry. Shown is the percentage of infected cells as measured by E protein staining (4G2+) (H) and viral RNA relative expression by RT-qPCR (I). One-way ANOVA was used to analyse these experiments. EV: empty vector. Error bars represent mean ± SD. Results are representative of two independent experiments. (J and K) A549 WT and ISG15 KO were infected with 20 ZIKV PFUs. At 36 hpi cells were fixed, permeabilized and stained for the flavivirus E protein. ZIKV relative foci area (J) and number of infected cells per foci (I), quantified by ImageJ software and analysed using unpaired t test with Welch’s correction. Images were acquired with an Olympus IX83 inverted microscope. Error bars represent mean ± SD. Results are representative of three independent experiments. Statistical analyses were performed using Prism 8 (GraphPad Software). p values *<0.05; **<0.01; ***<0.001.

Finally, lack of ISG15 expression led to an increase in foci size in cells infected with ZIKV, another flavivirus (Figure 2J and K) but not with HSV-1 (Figure S2A and B) or VSV (Figure S2C and D). These results are in line with Speer and colleagues’ data where cells isolated from humans deficient for ISG15 do not have enhanced susceptibility to HSV-1 or VSV [29]. Altogether these results suggest a specific role for ISG15 in the regulation of flavivirus replication and dissemination.

### ISGylation deficiency does not affect DV spread

ISG15 is an IFN-I-inducible ubiquitin-like molecule and can be conjugated to target proteins by HERC5, an ISG15 ligase also induced by IFN [23, 41]. Of note, ISGylation of host or viral proteins was reported to inhibit replication of several viruses such as influenza (IAV) [42], human cytomegalovirus (HCMV) [43] and human respiratory syncytial virus (RSV) [44]. Moreover, DV proteins were also shown to be ISGylated [45]. Considering that HERC5 is the only ISG15 ligase expressed in the dataset analysed, we generated HERC5-deficient A549 cells using CRISPR/Cas9 (Figure S3A) to determine whether ISGylation could be involved in DV restriction. Accordingly, A549 HERC5 null cells were not able to perform ISGylation after IFN-I treatment (Figure 3A). In contrast to ISG15 null cells, we did not observe differences between WT and HERC5-deficient cells when we evaluated both foci area (Figure 3B) and number of infected cells per foci following infection with DV (Figure 3C). Therefore, ISGylation is not sufficient to inhibit DV replication in this model.

**Figure 3.**
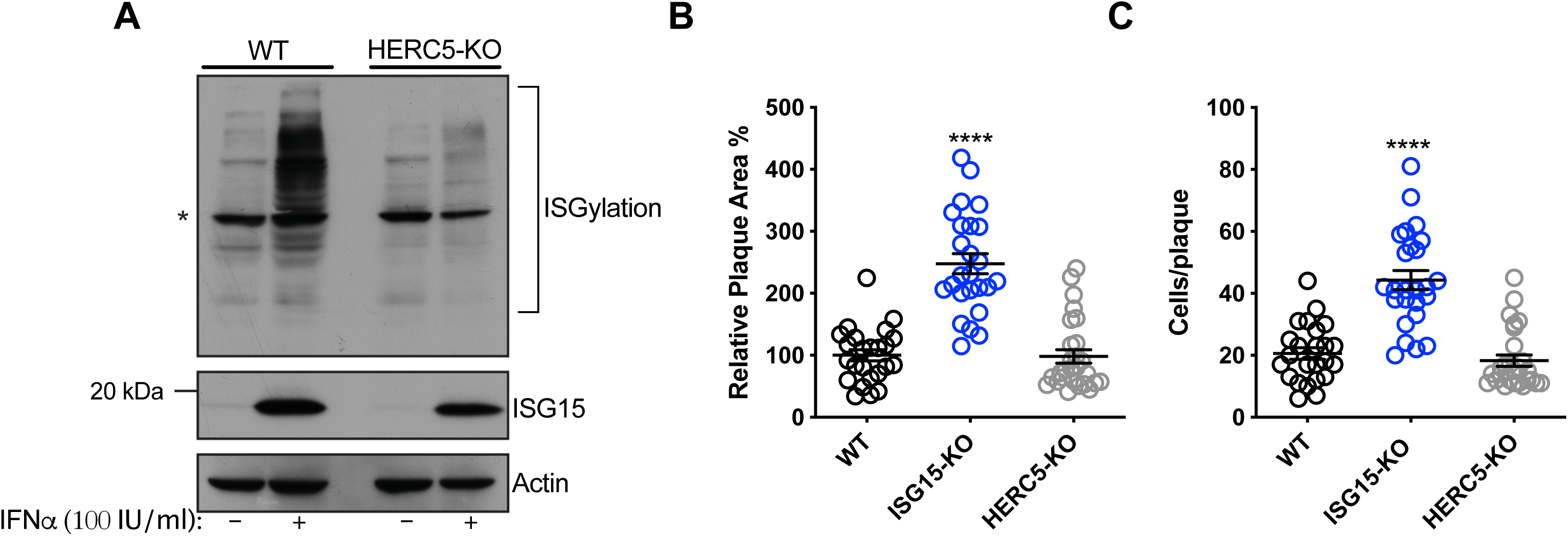
ISGylation is not sufficient to restrict DV spread. (A) ISGylation profile of A549 WT and HERC5 KO cells by Western blot. Cells were primed with IFNα2b (100 IU/ml) for 24 h and cell lysates were analysed with an ISG15 antibody. (*) indicates antibody unspecific band. (B and C) A549 cells were infected with 20 DV PFUs. At 36 hpi cells were fixed, permeabilized and stained for the flavivirus E protein. DV relative foci area (B) and the number of infected cells per foci (C) quantified by ImageJ software and analysed by one-way ANOVA. Images were acquired with an Olympus IX83 inverted microscope. Error bars represent mean ± SD. Results are representative of three or more independent experiments. Statistical analyses were performed using Prism 8 (GraphPad Software). p values ****<0.0001.

### ISG15 is necessary for autocrine IFNAR1-mediated control of DV replication

ISG15 and its binding partner USP18 are crucial for IFN-I pathway down-regulation, which is pivotal for infection control and immune-regulation in humans [28, 29, 46]. The A549 ISG15-KO cell line exhibited a lower expression of USP18 and sustained ISG expression, as exemplified by IFIT3, after IFN-I stimulation (Figure 4A), reproducing the phenotype observed in cells isolated from humans lacking ISG15 [29].

**Figure 4.**
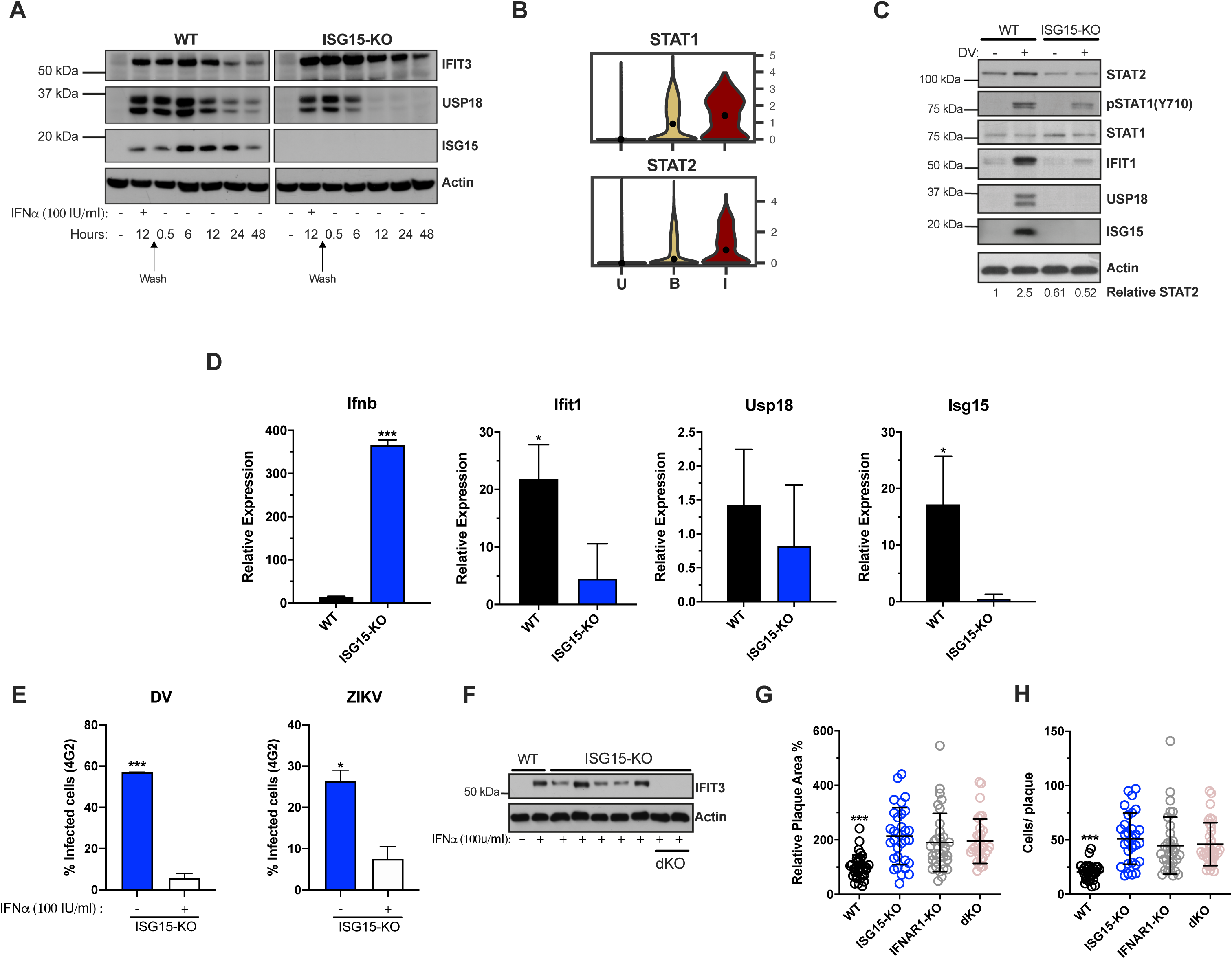
ISG15 is necessary for autocrine IFNAR1-mediated control of DV replication. (A) A549 cells were primed with IFNα2b (100 IU/ml) for 12 h, washed three times with DPBS and allowed to rest. Cells were harvested at the indicated time point and cell lysates were analysed by Western blot with the corresponding antibodies. Results are representative of two independent experiments. (B) Violin plot representing the expression of STAT1 and STAT2 in uninfected [U], bystander [B] and infected [I] PMBC. (C and D) A549 cells were infected with 20 DV PFU. At 36 hpi, cells were harvested, cell lysates were analysed by immunoblotting (C) and the indicated mRNA transcripts were quantified by RT-qPCR (D). Error bars represent mean ± SD. Results are representative of two independent experiments. Data was analysed by unpaired t test. (E) A549 ISG15 KO cells were primed with IFNα2b (100 IU/ml) for 12 h, washed three times with DPBS and allowed to rest 12 h before infection with DV or ZIKV at an MOI of 0.1. Shown is the percentage of cells infected at 36 hpi, as measured by E protein staining (4G2+). Error bars represent mean ± SD. Results are representative of three independent experiments. Data was analysed using unpaired t test. (F) Clonal isolates of A549 cells were immunoblotted for IFIT3 after 24 h treatment with IFNα2b (100 IU/ml) (+, control IFIT3-proficient cells; 1–9, different clonal isolates after limiting dilution). (G and H) A549 cells were infected with 20 DV PFUs. At 36 hpi cells were fixed, permeabilized and stained for the flavivirus E protein. DV relative foci area (G) and number of infected cells per foci (H) quantified by ImageJ software and analysed by one-way ANOVA. Images were acquired with an Olympus IX83 inverted microscope. Error bars represent mean ± SD. Results are representative of three independent experiments. Statistical analyses were performed using Prism 8 (GraphPad Software). p values *<0.05; ***<0.001.

While our single-cell RNA-seq analysis showed that both ISG15 and USP18 are upregulated in PMBCs containing DV RNA, there was an asynchrony of ISGs being expressed in infected versus bystander cells, with a greater spectrum present in the latter (Figure 1C). This is expected as flavivirus’ control of IFN-I signalling occurs mainly via its intracellular non-structural proteins [47, 48] and few ISGs can be directly induced by IRF3 [49–51].

Of interest, STAT2 mRNA is expressed at higher levels in DV RNA-containing cells when compared to bystanders and uninfected cells (Figure 1C and 4B). Hence, we hypothesised that ISG15 interference with DV replication is associated with autocrine IFN-I signalling in infected cells. Confirming the results from PMBCs and HuH7 cells, ISG15 mRNA was induced in A549 cells during DV infection (Figure 4D). As expected, this expression was abolished in knockout cells. Therefore, we measured the activation of the IFN-I pathway in WT and ISG15-KO cells infected with DV. Despite ISG15-knockout cells having full machinery to control viral infection as well as hyper-responsiveness to exogenous IFNα (Figure 4A) [28, 29], they did not respond properly to DV infection. Infected ISG15-deficient cells showed reduced STAT1 phosphorylation and STAT2 and IFIT1 expression when compared to WT cells. As expected, A549 lacking ISG15 had no detectable USP18 (Figure 4C). Infected knockout cells induced IFNβ mRNA at higher levels than the WT but had less IFIT1 mRNA (Figure 4D) indicating that DV infection impairs the response at both mRNA and protein levels downstream of IFN-I induction. Of note, ISG15 KO cells are still able to control DV and ZIKV infection when previously stimulated with IFNα (Figure E).

Taken together, our results suggest that ISG15 inhibits DV infection by regulating early events of the IFN-I receptor activation. To test this, we constructed an ISG15/IFNAR1 double knockout cell line based on the ISG15-KO background (Figure 4F). Cells were infected with DV and foci area and number of infected cells per foci was quantified. In parallel, we performed this experiment in IFNAR1 knockout cells, previously generated by our group [52, 53]. The same phenotype was observed in all three cell lines, with no additive or synergistic effects in the double knockout (Figures 4G and H), which places intracellular ISG15 downstream of IFNAR1 signalling in the control of DV infection.

### ISG15 counteracts DV IFN-I evasion

Flaviviruses are known to counteract IFN-I signalling by inducing the degradation of STAT2, a key protein in the interferon signal transduction pathway [11, 14, 54]. As our previous results suggest that ISG15’s role during DV infection is downstream of IFNAR engagement (Figure 4), we evaluated DV-mediated STAT2 degradation in the absence of ISG15. In agreement with others [55], STAT2 degradation in the A549 cell line occurs rapidly after infection. Despite showing sustained activation of IFN-I signalling (Figure 4A), ISG15-KO cells showed pronounced STAT2 degradation when infected with DV (Figure 5A and E-upper panel). As infected IFN-secreting cells are able to induce an antiviral state in neighbouring-bystander cells by inducing the expression of ISGs [56], we evaluated in which cell population, infected and/or bystander, ISG15 impacted DV infection. DV foci were co-stained for flavivirus E protein and IFIT3 and confocal microscopy was performed. Both infected and bystander cells were able to respond to infection, producing IFIT3 (Figure 5D, top panel). However, IFIT3 expression in the foci context was largely impaired and restricted to infected cells in the absence of ISG15 (Figures 5B, C and D). This data suggests that restriction of DV replication and dissemination is achieved by both autocrine and paracrine IFN-I response amplification and is dependent on ISG15 expression. To further investigate this, we sorted DV prM protein positive and negative cells (Figure S4A and B), therefore enabling us to evaluate the impact of infection in virus-containing and bystander cells, respectively. Strikingly, DV-positive cells had a lower expression of STAT2, IFIT3 and USP18 when compared to bystander cells; a phenotype that was markedly enhanced in ISG15-deficient cells (Figure 5E, *bottom panel*). While also confirming DV inhibition of IFNAR signalling, our results further suggest that ISG15 function is targeted for successful viral infection.

**Figure 5.**
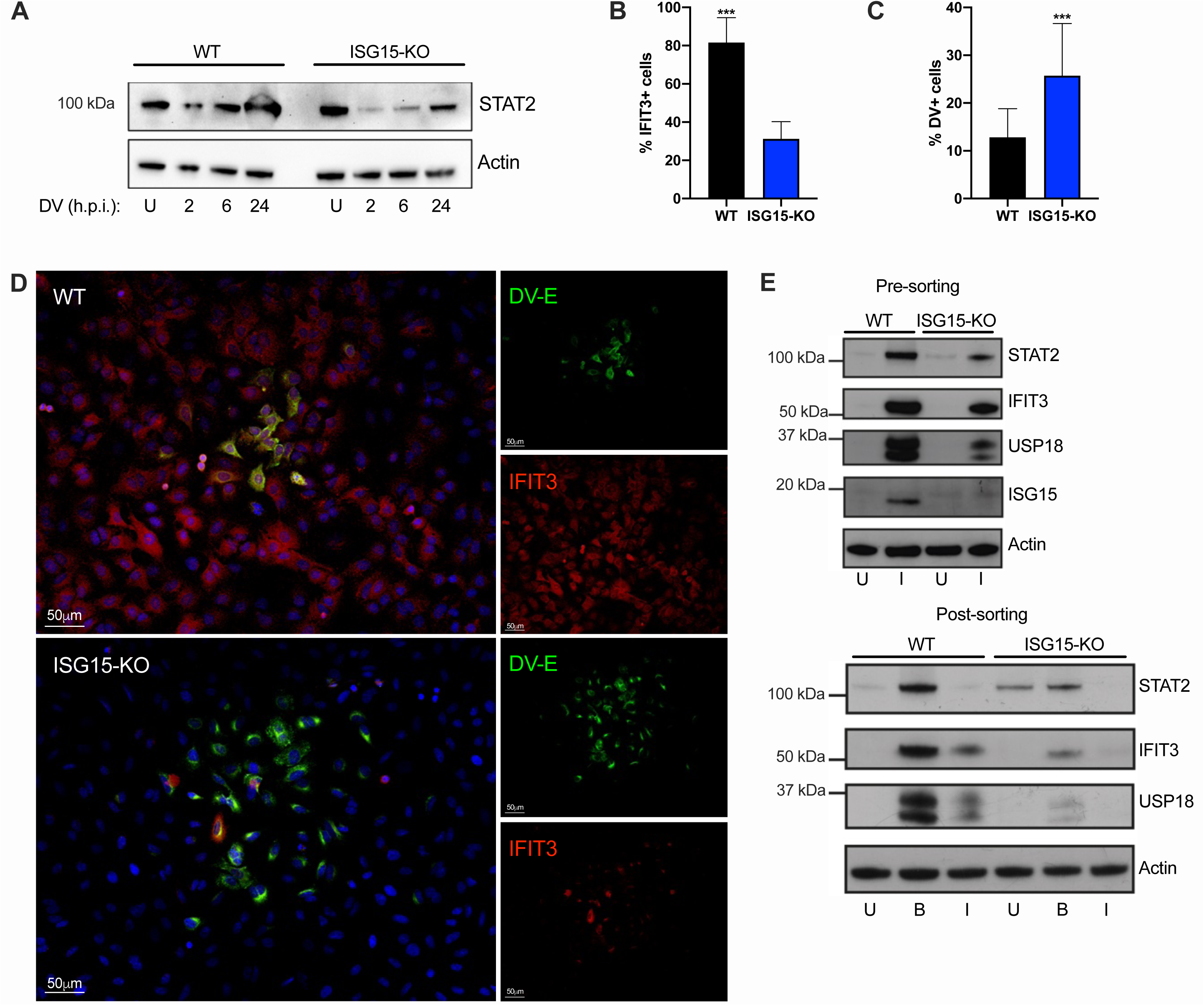
ISG15 counteracts DV IFN-I evasion. (A) A549 WT and ISG15 KO were infected with 20 DV PFU. Cells were harvested at the indicated time points after infection (hpi) and cell lysates were analysed by Western blot using STAT2 and Actin antibodies. (B, C and D) A549 WT and ISG15 KO immunofluorescence assay (IFA) 36 hpi for cellular IFIT3 and flavivirus E protein expression (D). Percentage of IFIT3 (B) and DV (C) positive cells per foci were quantified by ImageJ software and analysed using unpaired t test with Welch’s correction when appropriate. Displayed images were acquired with a Leica DMI6000 B microscope. (E) A549 cells were infected with DV at MOI 0.01. At 36 hpi, cells were fixed, permeabilized and stained for flavivirus E protein. Cell lysates were analysed by Western blot with the indicated antibodies before (upper panel) and after (bottom panel) cells were sorted by fluorescence-activated cell sorting (FACS) based on E protein expression. U: uninfected. B: bystander. I: infected Error bars represent mean ± SD. Results are representative of two independent experiments. Statistical analyses were performed using Prism 8 (GraphPad Software). p values ***<0.001.

### USP18 expression displaces NS5 from STAT2 and overcomes ISG15 deficiency

STAT2 degradation mediated by all DV serotypes and ZIKV is largely dependent on NS5 [11, 14], the virus RdRp and methyltransferase. NS5 was shown to bind to the N-terminus of human STAT2 [11, 15]. Interestingly, USP18 interacts with STAT2 via its coiled-coil and DNA binding domains. This causes negative regulation of IFNAR signalling by displacing JAK1 from chain 2 of the receptor [25]. As shown here (Figures 4A, C and 5E) and elsewhere [28, 29], the absence of ISG15 results in USP18’s destabilisation. We therefore examined if NS5, STAT2 and USP18 are part of the same complex. In HEK293 cells lacking ISG15 (Figure S3B, C and D) and primed with IFNα for 18 hours, both endogenous STAT2 and overexpressed USP18, immuno-precipitated with ZIKV’s NS5 (Figure 6A, lane 2). This interaction was enhanced when a protease-deficient high-expressing USP18 mutant (C64A) was used (Figure 6A, lane 3). We then hypothesised that the interaction of USP18 with STAT2 competes with NS5 binding, which in turn could result in a lower efficiency of virus evasion mechanism. To test this, we performed an STAT2 (V5) pull-down in ISG15-deficient HEK293 cells transfected with NS5, USP18 or a combination of both, followed by treatment with IFNα overnight. Increasing concentrations of USP18 (Figure 6B lanes 3, 4 and 5) show a concentration-dependent displacement of NS5 from STAT2, suggesting that both proteins indeed compete for the same region of the protein.

**Figure 6.**
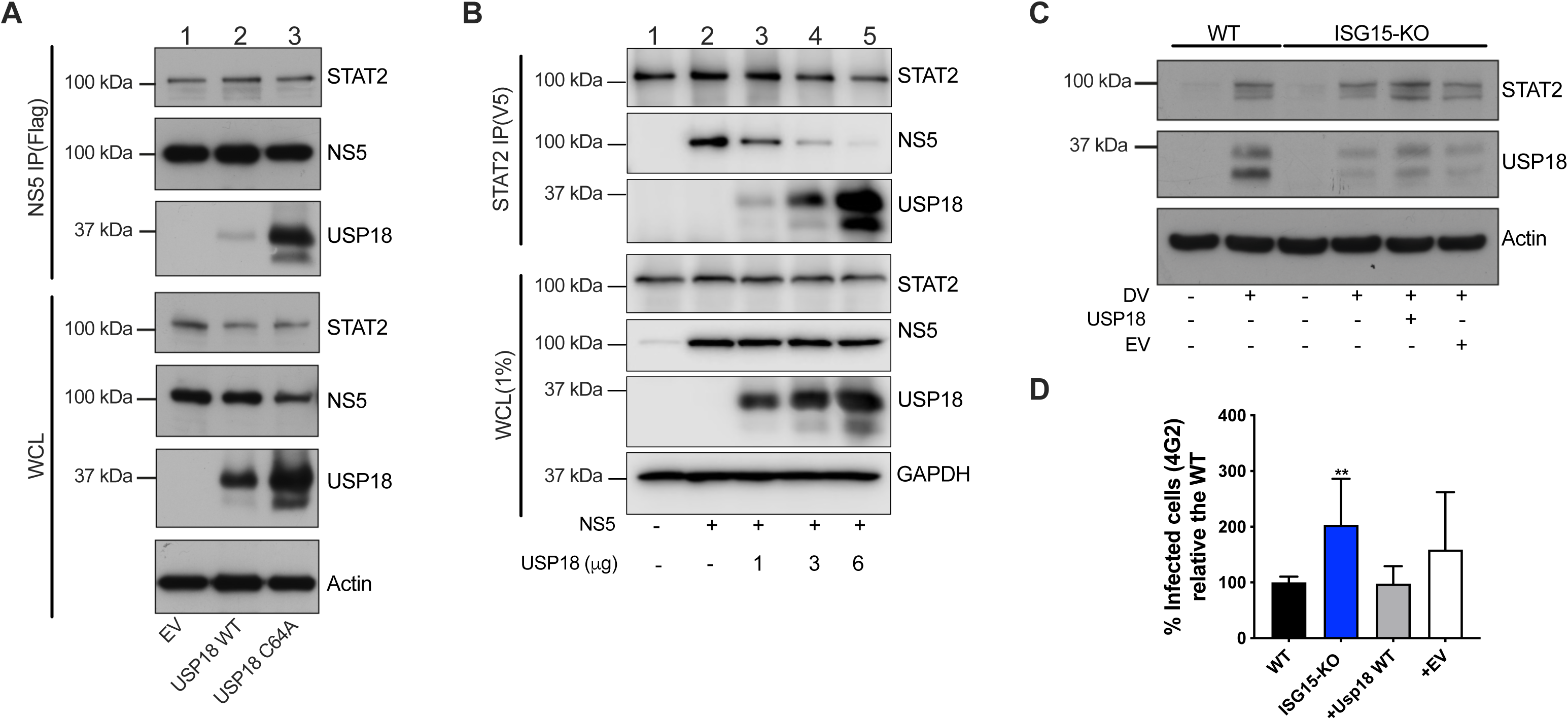
USP18 expression displaces NS5 from STAT2 and overcomes ISG15 deficiency. (A) Flag-tag immunoprecipitation (IP) assay and Western blot analysis of HEK293 ISG15 KO cells transfected with ZIKV NS5-FLAG, human USP18 WT, human USP18 C64A mutant or the empty vector (pcDNA3.1) plasmids, followed by IFNα2b (100 IU/ml) priming for 18 h. WCL, whole cell lysate. Results are representative of three independent experiments. (B) STAT2(V5) IP assay and Western blot analysis of HEK293 ISG15 KO cells transfected with the indicated plasmids, followed by IFNAα2b (100 IU/ml) priming for 18 h. Results are representative of three independent experiments. (C and D) Complementation with USP18 in A549 ISG15 KO cells. Cells were stably transfected with human USP18 or the empty vector and infected with DV at an MOI of 0.01. At 36 hpi, cells were harvested and cell lysates were analysed by Western blot with the corresponding antibodies (C). Percentage of cells infected as measured by E protein staining (D). Error bars represent mean ± SD. Results are representative of two independent experiments. Statistical analyses were conducted using Mann-Whitney’s test in Prism 8 (GraphPad Software. p value **<0.01 (D). EV: empty vector.

Thus, we evaluated whether the presence of USP18 could restore STAT2 expression in ISG15 knockout cells during DV infection. Overexpression of USP18 led to an increase of STAT2 levels similar to ones seen in wild type cells (Figure 6C). Importantly, USP18 reconstitution in ISG15-deficient cells was also able to reduce virus replication (Figure 6D), suggesting that reestablishment of the IFN-I signalling was sufficient to recover the WT phenotype in ISG15 knockout cells.

Together, the results presented here reveal that human ISG15 restricts DV and ZIKV replication via its ability to stabilise USP18 and regulate the type 1 IFN signalling pathway.

## DISCUSSION

A successful infection is dependent on the virus replication machinery and its ability to evade host immunity. One of the first lines of defence a virus has to overcome is a plethora of antiviral genes induced by IFN-I [57].

ISG15 is induced early during infection [33, 58] and has been shown to have a viral restriction role in several infection models. Most of those have been reported to be a consequence of ISGylation [42–44, 59], even though this process was suggested to be both inefficient and unspecific [59]. Here we show that during the IFN-I response elicited throughout DV infection, direct viral protein ISGylation is redundant for antiviral immunity; rather, ISG15’s ability to stabilise USP18 prevents NS5-mediated STAT2 degradation, thus leading to a more effective interferon response that culminates in DV and ZIKV restriction.

Many of ISG15’s functions have been shown to be immunomodulatory. For instance, ISGylation stabilises IRF3 by occluding its ubiquitylation sites [60], negatively regulates RIG-I [61] and activates PKR [62]. Secreted ISG15 functions as a cytokine [16, 63], leading to the production of IFNγ and IL-10 in human cells, crucial to the control of pathogens such as *Mycobacterium tuberculosis* [17, 19]. Moreover, free intracellular ISG15 is essential for USP18 stability (Figures 4C, 5E and [28, 29]) and its absence leads to severe interferonopathy in humans [28]. Also there are reports of ISG15 having an antiviral function independent of Ube1L during Chikungunya virus infection in mice [28, 64], where free ISG15 contributes to infection control by blunting potentially pathologic levels of cytokine effectors. Considering this range of functions, it is expected that different pathogens might interact with this pathway in different ways, according to its co-evolutionary history.

Of note, NK, NKT and monocytes were the PBMC populations with higher upregulation of ISG15 mRNA in single-cell gene expression studies. These have been shown to be the major producers and/or targets of free extracellular ISG15 in other contexts [17–19, 63]. The influence of extracellular ISG15 during viral infections should be further explored in the future.

Interestingly, humans lacking ISG15 do not have increased susceptibility to common viral infections, such as influenza and HSV-1. The explanation for this, as suggested elsewhere, may lie in the sustained IFN-I response of their cells creating a hostile environment for virus growth [28, 29]. Here, we reveal that ISG15 restricts DV and ZIKV replication indirectly by stabilising USP18 and thereby disrupting NS5-STAT2 interaction: ISG15 promotes competition for a niche exploited by such viruses. We also demonstrate that USP18, STAT2 and NS5 co-immunoprecipitate, suggesting a very narrow window of opportunity that NS5 has to degrade STAT2. As USP18/ ISG15 interaction is reported to down-regulate IFN-I signalling in humans but not mice [29], it is tempting to speculate that NS5 interaction with STAT2, a major flavivirus immune evasion mechanism and also restricted to humans [15], was shaped by the USP18/ISG15 interaction.

Our results suggest an unexpected mechanism by which ISG15 can exert an antiviral function distinct from the debilitating effects of its conjugation to viral proteins. The key role of IFN-I in viral infections might lead to the perception of ISGs having a necessarily direct antiviral function, a paradigm that is recently being reassessed, with a range of ISGs being implicated in infection-independent functions [65, 66]. Here we provide mechanistic insight of the arms race between ISG15, USP18 and NS5 which suggests that protein/protein dynamics adjacent to IFNAR are a key determinant for the outcome of flavivirus infection.

## ACKNOWLEDGMENTS

This work was supported by Comissão de Aperfeiçoamento de Pessoal de Nível Superior (CAPES) Computational Biology programme (23038.010048/2013-27), Conselho Nacional de Desenvolvimento Científico e Tecnológico (CNPq) (311406/2017-3 and 407609/2018-0), INCT Vacinas/CNPq and the Academy of Medical Sciences/U.K. (NAF004/1005) to D.S.M. Howard Hughes Medical Institute – Early Career Scientist to A.B. (55007412), Wellcome Trust and Royal Society Sir Henry Dale Fellowship (202471/Z/16/Z) to T.R.S. Wellcome Trust PhD Studentship (105389/Z/14/Z) to T.J.S. CAPES scholarship (88887.195782/2018-00) to E.L.R. CNPq scholarship (205096/2018-2) to J.B.

We thank Dr. Sandra Pellegrini and Dr. Carsten Münk for kindly providing human USP18 plasmids. We thank Dr. Brian Ferguson for critical reading of the manuscript.

## AUTHOR CONTRIBUTIONS

Conceptualization, C.E.E., A.B. and D.S.M.; Methodology, C.E.E., E.L.R and D.S.M.; Software, E.L.R; Formal Analysis, C.E.E., E.L.R and E.G.K., Investigation, C.E.E., A.A.S., D.O.P, G.M. and JB; Resources, C.E.E., A.A.S, P.F.S., Z.G, T.J.S., T.F. and T.R.S. Data Curation, E.L.R. and E.G.K., Writing-Original Draft, C.E.E. and D.S.M., Writing-Review and editing, C.E.E, E.L.R, D.O.P., E.G.K., P.F.S., J.B., T.J.S., T.F., T.R.S, A.B., and D.S.M., Visualization, C.E.E., E.L.R. and D.S.M., Supervision, D.S.M., Project Administration and Funding Acquisition, D.S.M.

## DECLARATION OF INTERESTS

The authors declare no competing interests.

## LEAD CONTACT AND MATERIAL AVAILABILITY

Further information and requests for resources and reagents should be directed to and will be fulfilled by the Lead Contact, Daniel Santos Mansur

## MATERIAL AND METHODS

### Cells and viruses

Mammalian cell lines were maintained at 37 °C under the conditions of a humidified atmosphere and 5% CO2. The human alveolar adenocarcinoma-derived A549 cells, human embryonic kidney HEK293 and the African green monkey kidney-derived Vero cells were cultured in Dulbecco’s modified Eagle’s medium F-12 (DMEM F12) (Gibco) supplemented with 5% foetal bovine serum (FBS) (Gibco) and streptomycin/penicillin (100 U/ml) (Gibco). The *Aedes albopictus* mosquito-derived cell line C6/36 was maintained at 28°C in a BOD in Leibovitz’s L-15 medium (L-15) (Gibco) supplemented with 10% FBS (Gibco), 0.26% tryptose phosphate broth (Sigma) and 50 µg/ml gentamicin (Sigma). A549 ISG15 KO and IFNAR1-KO have been described elsewhere [17, 52, 53]. All cells were negative for mycoplasma.

Dengue virus serotype 4 (DENV-4 TVP/360 – GenBank accession number: KU513442) and Zika virus (ZV BR 2015/15261 - GenBank accession number: MF073358) stocks were propagated in C6/36 cells and titrated in Vero cells. Vesicular stomatitis virus-green fluorescent protein (VSVeGFP) (Indiana strain, Marques-JT, Plos Pathogens 2013) and Herpes simplex virus-1-green fluorescent protein (HSV-1eGFP) (SC16) viruses were propagated and titrated in Vero cells. HSV-1eGFP was a kind gift of Professor Stacey Efstathiou.

### Single-cell RNA sequencing analysis

Processed, publicly available single-cell RNA-seq data are available through the GEO accession numbers GSE116672 and GSE110496. We downloaded processed single-cell data and metadata from the supplementary information from the respective publications [31, 32].

Then, we used CellRouter to analyse these datasets. To perform the tSNE analysis using single-cell data generated by Zanini 2018 [32], we set the parameters num.pcs=10, seed=1 and max_iter=1000 in the computeTSNE function. Next, we identified genes preferentially expressed in Uninfected, Bystander and Infected cells using a cutoff for the log2 fold change of 0.25. We used a custom script to obtain all genes containing the keywords “type I interferon” in the Gene Ontology Biological Processes (package versions: org.Hs.eg.db_3.10.0, GO.db_3.10.0). Next, we took the overlap of type I interferon genes with the genes preferentially in each condition reported above. The remaining analyses were focused on these genes.

To perform the tSNE analysis using the single-cell data generated by Zanini 2018 [31], we set the parameters num.pcs=20, seed=1 and max_iter=1000 in the computeTSNE function. We used a strategy similar to the one described above to identify genes differentially expressed in each condition but used a cutoff of 0.15 for this dataset. The parameter num.pcs was determined using the elbow approach, as described in the CellRouter tutorial at https://github.com/edroaldo/cellrouter.

### CRISPR/Cas9-mediated gene editing

A549 WT and ISG15 KO cells were co-transfected with two Herc5 or Ifnar1-targeting gRNA CRISPR/Cas9-GFP plasmids, respectively. HEK293 WT cells were transfected with three Isg15-targeting gRNA CRISPR/Cas9-GFP plasmids (Table S1) (Horizon Cambridge, UK). After 72 h, cells were sorted by FACS (FACSMelody, BD) and single-cell derived clones were initially screened by PCR genotyping. Additionally, both HERC5 and ISG15/IFNAR1 (dKO) clones were functionally tested by assessing their ISGylation profile and expression of ISGs, respectively, after IFNα priming. Briefly, A549 WT and HERC5 and dKO clones were primed with IFNα2b (100 IU/ml) (PBL Assay Science) for 24 h and the expression of ISG15-conjugates and IFIT3 was analysed by Western blot. HEK293 cells were primed with IFNα2b (1000 IU/ml) for 8 h, total RNA was isolated and Isg15 mRNA expression was assessed by RT-qPCR.

### Viral infection

A549 cells were seeded one day prior to infection in appropriate multi-well plates. For foci assay, a viral inoculum containing 20 foci forming units (PFU) of the corresponding virus was added to each well, and virus adsorption was performed in DMEM supplemented with penicillin/streptomycin (100 U/ml) for 90 min at 37°C. Cells were washed with PBS to remove un-adsorbed virus, and maintained in DMEM 1.5% carboxymethylcellulose sodium (CMC) (Sigma). Alternatively, cells were infected with DV at the indicated MOI, as described above, and maintained in DMEM supplemented with 1% FBS.

### Titration

DV titration was performed by focus forming assay (FFA) in C6/36 cells. Briefly, C6/36 cells were seeded (1×10^5^ cells/well in 24 well plate) and after overnight incubation were infected with a 10-fold serial dilution of virus samples (cell culture supernatants) in L-15 with 0.26% tryptose and 25 μg/ml of gentamicin. After 90 minutes, the inoculum was removed and a CMC overlay media (L-15 media with 5% FCS, 0.26% tryptose, 25 μg/ml of gentamicin and 1.6% of CMC) was added and the plates were incubated for 7 days at 28°C. After incubation, cells were washed, fixed with 3% paraformaldehyde (PFA) (Sigma-Aldrich) and permeabilized with 0.5% triton X100 (Sigma-Aldrich). After washing, cells were immunostained with mouse monoclonal anti flavivirus E protein antibody 4G2 (ATCC® HB-112™, dilution 1:100), followed by goat anti-mouse immunoglobulin conjugated to alkaline phosphatase (Promega S3721, dilution 1:7500). Focuses of infection were revealed using NBT/BCIP reagent (Promega), following the manufacturer’s instructions and the virus titer calculated as follow: media of focus number/inoculum volume x dilution. The results are expressed as FFU_C6/36_/ml (Gould et al., 1985).

VSVeGFP and HSV-1eGFP titrations were performed by foci assay. Briefly, Vero cells were seeded in 24-well plates 24 h prior to infection. Cell monolayers were washed with PBS and inoculated with 0.3 ml of serial 10-fold dilutions of the virus in duplicates. After 90 min adsorption at 37°C, each well was re-suspended in DMEM 1.5% CMC. At 48 hpi, cells were fixed with 3% PFA (Sigma) for 30 min, washed 3 times with PBS and stained with 1% crystal violet (Sigma) for 30 min at room temperature. Virus yield was calculated and expressed as foci forming units per ml (PFU/ml).

### Immunofluorescence

At 24 hpi and 18 hpi, respectively, HSVeGFP and VSVeGFP infected cells were fixed with 3% PFA for 20 min at room temperature, permeabilized with 0.5% Triton X100 in PBS for 4 min and stained with DAPI counterstain (Molecular Probes). At 36 hpi, DV and ZIKV infected cells were fixed, permeabilized and stained with mouse monoclonal 4G2 antibody (10 µg/ml dilution 1:100), followed by incubation with Alexa Fluor 488 rabbit anti-mouse IgG (HL+LL, Life Technologies, dilution 1:500), and DAPI counterstain. Images were acquired with an Olympus IX83 inverted microscope. Briefly, virus focis, determined by eGFP or flavivirus E protein expression, were delimited; images were converted to 16-bit and processed to be analysed with the ImageJ Software Cell Counter Plugin (W. S. Rasband, ImageJ, US National Institutes of Health, Bethesda, MD; http://rsb.info.nih.gov/ij/, 1997–2006).

For confocal analysis, A549 cells grown on glass coverslips were mock infected or infected with 10 DV PFU. At 36 hpi, cells were fixed and permeabilized. Following washes with PBS, cells were stained with mouse monoclonal anti-E protein (4G2, dilution 1:100) and rabbit polyclonal anti-IFIT3 (Proteintech, 15201-1-AP, dilution 1:200) for 1 h at room temperature. The cells were washed with PBS and stained with secondary antibodies Alexa Fluor 488 goat anti-mouse IgG (HL+LL, Life Technologies, dilution 1:500) and Alexa 568 goat anti-rabbit IgG (H+L, Life Technologies, dilution 1:500) and DAPI counterstain (Molecular Probes). Cells were washed and coverslips mounted using Prolong antifade reagent (Invitrogen). Z-stack and max intensity projection images were generated with a Leica DMI6000 B confocal microscope and Leica Application Suite X software for image analysis (Leica Microsystems).

### Flow cytometry and cell sorting

Cells were fixed with 2% PFA in PBS, washed twice with PBS and permeabilized with 0.5% saponin in 1% BSA in PBS. Anti-flavivirus E protein mAb 4G2 was conjugated to Alexa Fluor 488 5-SDP (Life Technologies) following the manufacturer’s instructions. Cells were incubated with FITC-conjugated 4G2 (dilution 1:1000) in permeabilization buffer for 40 minutes at room temperature, washed once and resuspended in FACS buffer. The cell suspensions were analysed by flow cytometry on a FACSVerse instrument (BD Biosciences) and analysed using FlowJo V10 software (BD). Cell sorting experiments were performed on a FACSMelody cell sorter (BD Biosciences).

### RT-qPCR

vRNA was isolated by using QIAamp viral RNA Mini Kit (Qiagen). Intracellular total RNA was isolated with TRIZOL (Thermo Life) following manufacturer’s instructions. A total of 1 µg was reverse transcribed using the High capacity cDNA reverse transcription kit (Applied Biosystems). Quantitative PCRs (qPCRs) were performed with GoTaq ® qPCR Master mix (Promega) following the standard cycling conditions suggested by the manufacturer in a StepOnePlus real time PCR system (Applied Biosystems). VSVeGFP and 18S mRNA were used as control housekeeping genes. Amounts of DV or ISG mRNA were calculated by using the ΔΔCT method. Primers specific to the mRNA analysed are listed in Table S1.

### Western blot

Human antibodies used for immunoblot were as follows: mouse mAb to β-actin (Abcam, ab6276, dilution 1:4000), rabbit mAb to IFIT1 (Abcam, ab137632, dilution 1:1000), mouse mAb to pSTAT1(Y701) [M135] (Abcam, ab29045, dilution 1:1000), mouse mAb to STAT1 (Abcam, ab3987, dilution 1:1000), rabbit pAb to IFIT3 (ProteinTech, 15201-1-AP, dilution 1:1000), rabbit mAb to USP18 [D4E7] (Cell Signalling Technologies, 4813, dilution 1:1000), rabbit mAb to STAT2 [D9J7L] (Cell Signalling Technologies, 72604, dilution 1:1000), mouse mAb to ISG15 (R&D System, MAB4845, dilution 1:1000), home-made mouse mAb anti-GFP [67, 68] (dilution 1:1000) and mouse mAb to FLAG tag (Sigma, F3165, dilution 1:500).

Cells were treated as indicated, wash two times with ice-cold PBS and then lysed in RIPA buffer [50 mM tris-HCl (pH 7.5), 150 mM NaCl, 1% Triton X100, 0.5% sodium deoxycholate, 0.1% SDS, 5 mM EDTA] supplemented with 1x protease inhibitors (Mini Protease Inhibitor Tablets, Roche). Total protein concentration was determined by BSA Protein Assay Kit (Thermo Life). For western blotting, 20 µg of total protein were prepared in dithiothreitol-containing Laemmli sample buffer, separated and transferred to a nitrocellulose blotting membrane (GE Healthcare Amersham). After transfer, the membrane was blocked with 5% nonfat dry milk in TBS 0.1% Tween 20 (TBST) for 1 h at room temperature. Membrane was incubated with primary Abs diluted in 2% BSA in TBST at 4°C with gentle shaking overnight. Membrane was washed three times with TBST and then incubated with the appropriate secondary HRP-linked antibody for 1 h at room temperature. Membranes were washed and covered with ECL developing solution (Pierce^TM^ ECL WB substrate, Thermo Fisher Scientific).

### Plasmids and transfections

Herc5 and Ifnar1 sgRNA/Cas9/GFP plasmids were provided by Horizon (Cambridge, UK). sgRNA sequences are described in Supplementary Table 1. Expression plasmid for ISG15 was described elsewhere [17]. Pmax^TM^ GFP expression vector was acquired from Lonza (cat numb #D-00061). Expression plasmid for ZIKV NS5 was generated by amplifying the NS5 coding sequence (amino acids 2521-3423 in the polyprotein) from a previously described plasmid-based ZIKV reverse genetic system [69] using primers containing an N-terminal FLAG tag and inserted into pcDNA3.1. Expression plasmids for human USP18 WT and USP18 C64A mutants were kindly provided by Dr Carsten Münk [70]. Transfections were performed with FuGene6 (Promega), following the manufacturer’s instructions. Stably transfected cells were selected with geneticin (500 µg/ml) (Sigma).

### Immunoprecipitation

For Stat2 co-immunoprecipitation assays, cells were lysed in RIPA buffer composed of 20 mM Tris-HCl pH 7.4, 150 mM NaCl,10 mM CaCl2, 0.1% v/v Triton X-100, 10% v/v glycerol and complete protease inhibitor cocktail (Mini Protease Inhibitor Tablets, Roche). The supernatant was separated by centrifugation at 12.000 g at 4 °C for 10 min and incubated with STAT2 antibody (2 µg) (Santa Cruz Biotechnology, 514193) for 3 h at 4°C with gentle shaking. Complexes were precipitated with protein A/G Plus-agarose (Santa Cruz Biotechnology), washed with TBS and resuspended in SDS sample buffer. Immunoprecipitates were subjected to SDS-PAGE and western blotting, as described above. FLAG-tagged proteins were immunoprecipitated with anti-FLAG M2-agarose (Sigma), following the manufacturer’s instructions.

All assays were performed three times and representative blots are presented.

## QUANTIFICATION AND STATISTICAL ANALYSIS

Details concerning the statistical analysis methods are provided in each figure legend. Briefly, all data were analysed using GraphPad Prism 8 software and were shown as mean and the standard deviation (SD). Statistical significance was determined by Welch’s t test or one-way ANOVA for foci size analyses, unpaired t test for virus multiple-step growth curve, cellular mRNA quantification and percentage of cells infected. Statistical significance is indicated by ∗, p < 0.05; ∗∗, p < 0.01; ∗∗∗, p < 0.001.

## SUPPLEMENTAL INFORMATION

**Figure S1**

(A) tSNE was used to visualise the single-cell global transcriptome data. Blue dots represent uninfected cells, derived from healthy donors. Beige dots represent bystander cells and red dots represent infected cells, both derived from patients infected with DV.

(B) Differential expression of ISGs in HuH7 cells infected with DV. The Gene Ontology term “type one interferon” was used to filter the results from the single-cell RNA sequencing.

(C) Single cell ISG expression variability in uninfected, bystander and infected. ISGylation related genes are underlined. Blue bar represents uninfected cells. Beige bar represents bystander cells and red bar represents infected cells, both derived from cells exposed to DV.

(D) Violin plot representing the expression of ISGylation family members in uninfected [U], bystander [B] and infected [I] HuH7 cell line.

**Figure S2. ISG15 does not restrict HSV-1 and VSV spread**

(A and B) A549 cells were infected with 20 HSV-1eGFP PFU. At 24 hpi, cells were fixed and stained with DAPI counterstain. HSV-1eGFP relative foci area (A) and number of cells per foci (B). Images were acquired with an Olympus IX83 inverted microscope and quantified by ImageJ software.

(C and D) A549 cells were infected with 20 VSVeGFP PFU. At 18 hpi, cells were fixed and stained with DAPI counterstain. VSVeGFP relative foci area (C) and number of cells per foci

(D). Images were acquired with an Olympus IX83 inverted microscope and quantified by ImageJ software.

Error bars represent mean ± SD. Results are representative of three or more independent experiments. Statistical analyses were conducted using Mann-Whitney’s test in Prism 8 (GraphPad Software).

**Figure S3. Characterization of knockout cell lines**

(A) Agarose gel electrophoresis of A549 WT and HERC5 KO PCR products using primers surrounding CRISPR/Cas9 HERC5 sgRNA guides editing region. PCR product size: WT: 310 bp; HERC5 KO: ∼250 bp. (*) indicates PCR unspecific band. WT: A549 WT cell line; KO: A549 HERC5 KO; B: blank

(B) Agarose gel electrophoresis of HEK293 WT and ISG15 KO PCR products using primers surrounding CRISPR/Cas9 ISG15 sgRNA editing region. PCR product size: WT: 1016bp; ISG15 KO: ∼180bp. (*) indicates PCR unspecific band. WT: HEK293 WT cell line; KO: HEK293 ISG15 KO; B: blank

(C) HEK293 WT and ISG15 KO cells were stimulated with IFNα2b (1000 IU/ml) for 8 h. Cells were harvested and total RNA was isolated. Isg15 mRNA was analysed by RT-qPCR.

(D) HEK293 ISG15KO PCR product was cloned into pGEM vector and sequenced by Sanger method. Nucleotide sequence was aligned with the Isg15 reference sequence retrieved from GenBank (NM_005101) and translated into the primary amino acid sequence. (.) indicates the same sequence; (-) gap; (*) stop codon.

**Figure S4. Sorting of infected cells**

(A and B) A549 cells were infected with DV at MOI 0.01. Representative FACS profile and DV mRNA qPCR of WT (A) and ISG15-KO (B) cells sorted by flavivirus E protein expression (4G2, FITC-A axis).

B: bystander. I: infected

## Notes

### Competing Interest Statement

The authors have declared no competing interest.

### Summary of Updates

We rearrange the figures order

## REFERENCES

1. Mansur DS, Smith GL, Ferguson BJ. Intracellular sensing of viral DNA by the innate immune system. Microbes Infect 2014;16:1002–1012.

2. Gebhardt A, Laudenbach BT, Pichlmair A. Discrimination of Self and Non-Self Ribonucleic Acids. J Interferon Cytokine Res 2017;37:184–197.

3. Schoggins JW. Interferon-Stimulated Genes: What Do They All Do? Annu Rev Virol. 2019. DOI: 10.1146/annurev-virology-092818-015756.

4. García-Sastre A. Ten Strategies of Interferon Evasion by Viruses. Cell Host Microbe 2017;22:176–184.

5. Pierson TC, Diamond MS. The emergence of Zika virus and its new clinical syndromes. Nature 2018;560:573–581.

6. Barrett ADT. The reemergence of yellow fever. Science 2018;361:847–848.

7. Bhatt S, Gething PW, Brady OJ, Messina JP, Farlow AW, et al. The global distribution and burden of dengue. Nature 2013;496:504–507.

8. Shepard DS, Undurraga EA, Halasa YA, Stanaway JD. The global economic burden of dengue: a systematic analysis. Lancet Infect Dis 2016;16:935–941.

9. Miorin L, Maestre AM, Fernandez-Sesma A, García-Sastre A. Antagonism of type I interferon by flaviviruses. Biochem Biophys Res Commun 2017;492:587–596.

10. Ngono AE, Shresta S. Immune Response to Dengue and Zika. Annu Rev Immunol 2018;36:279–308.

11. Grant A, Ponia SS, Tripathi S, Balasubramaniam V, Miorin L, et al. Zika Virus Targets Human STAT2 to Inhibit Type I Interferon Signaling. Cell Host Microbe 2016;19:882–890.

12. Laurent-Rolle M, Morrison J, Rajsbaum R, Macleod JML, Pisanelli G, et al. The interferon signaling antagonist function of yellow fever virus NS5 protein is activated by type I interferon. Cell Host Microbe 2014;16:314–327.

13. Chen S, Wu Z, Wang M, Cheng A. Innate Immune Evasion Mediated by Flaviviridae Non-Structural Proteins. Viruses;9. Epub ahead of print 7 October 2017. DOI: 10.3390/v9100291.

14. Morrison J, Laurent-Rolle M, Maestre AM, Rajsbaum R, Pisanelli G, et al. Dengue virus co-opts UBR4 to degrade STAT2 and antagonize type I interferon signaling. PLoS Pathog 2013;9:e1003265.

15. Ashour J, Morrison J, Laurent-Rolle M, Belicha-Villanueva A, Plumlee CR, et al. Mouse STAT2 restricts early dengue virus replication. Cell Host Microbe 2010;8:410– 421.

16. Dos Santos PF, Mansur DS. Beyond ISGlylation: Functions of Free Intracellular and Extracellular ISG15. J Interferon Cytokine Res 2017;37:246–253.

17. Dos Santos PF, Van Weyenbergh J, Delgobo M, Oliveira Patricio D de, Ferguson BJ, et al. ISG15-Induced IL-10 Is a Novel Anti-Inflammatory Myeloid Axis Disrupted during Active Tuberculosis. J Immunol 2018;200:1434–1442.

18. Swaim CD, Scott AF, Canadeo LA, Huibregtse JM. Extracellular ISG15 Signals Cytokine Secretion through the LFA-1 Integrin Receptor. Mol Cell 2017;68:581–590.e5.

19. Bogunovic D, Byun M, Durfee LA, Abhyankar A, Sanal O, et al. Mycobacterial disease and impaired IFN-γ immunity in humans with inherited ISG15 deficiency. Science 2012;337:1684–1688.

20. Delgobo M, Daniel A G, Kozlova E, Rocha EL, Rodrigues-Luiz GF, et al. An evolutionary recent IFN-IL-6-CEBP axis is linked to monocyte expansion and tuberculosis severity in humans. DOI: 10.1101/514943.

21. Skaug B, Chen ZJ. Emerging role of ISG15 in antiviral immunity. Cell 2010;143:187– 190.

22. Dastur A, Beaudenon S, Kelley M, Krug RM, Huibregtse JM. Herc5, an interferon-induced HECT E3 enzyme, is required for conjugation of ISG15 in human cells. J Biol Chem 2006;281:4334–4338.

23. Wong JJY, Pung YF, Sze NS-K, Chin K-C. HERC5 is an IFN-induced HECT-type E3 protein ligase that mediates type I IFN-induced ISGylation of protein targets. Proc Natl Acad Sci U S A 2006;103:10735–10740.

24. Perng Y-C, Lenschow DJ. ISG15 in antiviral immunity and beyond. Nat Rev Microbiol 2018;16:423–439.

25. Arimoto K-I, Löchte S, Stoner SA, Burkart C, Zhang Y, et al. STAT2 is an essential adaptor in USP18-mediated suppression of type I interferon signaling. Nat Struct Mol Biol 2017;24:279–289.

26. Malakhov MP, Malakhova OA, Kim KI, Ritchie KJ, Zhang D-E. UBP43 (USP18) specifically removes ISG15 from conjugated proteins. J Biol Chem 2002;277:9976– 9981.

27. Malakhova O, Malakhov M, Hetherington C, Zhang D-E. Lipopolysaccharide activates the expression of ISG15-specific protease UBP43 via interferon regulatory factor 3. J Biol Chem 2002;277:14703–14711.

28. Zhang X, Bogunovic D, Payelle-Brogard B, Francois-Newton V, Speer SD, et al. Human intracellular ISG15 prevents interferon-α/β over-amplification and auto-inflammation. Nature 2015;517:89–93.

29. Speer SD, Li Z, Buta S, Payelle-Brogard B, Qian L, et al. ISG15 deficiency and increased viral resistance in humans but not mice. Nat Commun 2016;7:11496.

30. Lenschow DJ, Lai C, Frias-Staheli N, Giannakopoulos NV, Lutz A, et al. IFN-stimulated gene 15 functions as a critical antiviral molecule against influenza, herpes, and Sindbis viruses. Proc Natl Acad Sci U S A 2007;104:1371–1376.

31. Zanini F, Pu S-Y, Bekerman E, Einav S, Quake SR. Single-cell transcriptional dynamics of flavivirus infection. Elife;7. Epub ahead of print 16 February 2018. DOI: 10.7554/eLife.32942.

32. Zanini F, Robinson ML, Croote D, Sahoo MK, Sanz AM, et al. Virus-inclusive single-cell RNA sequencing reveals the molecular signature of progression to severe dengue. Proc Natl Acad Sci U S A 2018;115:E12363–E12369.

33. Ramos I, Smith G, Ruf-Zamojski F, Martinez-Romero C, Fribourg M, et al. Innate immune response to influenza virus at single-cell resolution in human epithelial cells revealed paracrine induction of interferon lambda 1. J Virol. Epub ahead of print 2 August 2019. DOI: 10.1128/JVI.00559-19.

34. Keskinen P, Nyqvist M, Sareneva T, Pirhonen J, Melén K, et al. Impaired antiviral response in human hepatoma cells. Virology 1999;263:364–375.

35. Schmid B, Rinas M, Ruggieri A, Acosta EG, Bartenschlager M, et al. Live Cell Analysis and Mathematical Modeling Identify Determinants of Attenuation of Dengue Virus 2’-O-Methylation Mutant. PLoS Pathog 2015;11:e1005345.

36. Roth H, Magg V, Uch F, Mutz P, Klein P, et al. Flavivirus Infection Uncouples Translation Suppression from Cellular Stress Responses. MBio;8. Epub ahead of print 10 January 2017. DOI: 10.1128/mBio.02150-16.

37. Wang K, Zou C, Wang X, Huang C, Feng T, et al. Interferon-stimulated TRIM69 interrupts dengue virus replication by ubiquitinating viral nonstructural protein 3. PLoS Pathog 2018;14:e1007287.

38. Whelan JN, Li Y, Silverman RH, Weiss SR. Zika Virus Production Is Resistant to RNase L Antiviral Activity. J Virol;93. Epub ahead of print 15 August 2019. DOI: 10.1128/JVI.00313-19.

39. Gullberg RC, Steel JJ, Pujari V, Rovnak J, Crick DC, et al. Stearoly-CoA desaturase 1 differentiates early and advanced dengue virus infections and determines virus particle infectivity. PLoS Pathog 2018;14:e1007261.

40. Zhang J, Lan Y, Li MY, Lamers MM, Fusade-Boyer M, et al. Flaviviruses Exploit the Lipid Droplet Protein AUP1 to Trigger Lipophagy and Drive Virus Production. Cell Host Microbe 2018;23:819–831.e5.

41. Zhao C, Denison C, Huibregtse JM, Gygi S, Krug RM. Human ISG15 conjugation targets both IFN-induced and constitutively expressed proteins functioning in diverse cellular pathways. Proc Natl Acad Sci U S A 2005;102:10200–10205.

42. Zhao C, Hsiang T-Y, Kuo R-L, Krug RM. ISG15 conjugation system targets the viral NS1 protein in influenza A virus-infected cells. Proc Natl Acad Sci U S A 2010;107:2253–2258.

43. Kim YJ, Kim ET, Kim Y-E, Lee MK, Kwon KM, et al. Consecutive Inhibition of ISG15 Expression and ISGylation by Cytomegalovirus Regulators. PLoS Pathog 2016;12:e1005850.

44. González-Sanz R, Mata M, Bermejo-Martín J, Álvarez A, Cortijo J, et al. ISG15 Is Upregulated in Respiratory Syncytial Virus Infection and Reduces Virus Growth through Protein ISGylation. J Virol 2016;90:3428–3438.

45. Hishiki T, Han Q ’en, Arimoto K-I, Shimotohno K, Igarashi T, et al. Interferon-mediated ISG15 conjugation restricts dengue virus 2 replication. Biochem Biophys Res Commun 2014;448:95–100.

46. François-Newton V, Magno de Freitas Almeida G, Payelle-Brogard B, Monneron D, Pichard-Garcia L, et al. USP18-based negative feedback control is induced by type I and type III interferons and specifically inactivates interferon α response. PLoS One 2011;6:e22200.

47. Dalrymple NA, Cimica V, Mackow ER. Dengue Virus NS Proteins Inhibit RIG-I/MAVS Signaling by Blocking TBK1/IRF3 Phosphorylation: Dengue Virus Serotype 1 NS4A Is a Unique Interferon-Regulating Virulence Determinant. MBio 2015;6:e00553–15.

48. Muñoz-Jordán JL, Laurent-Rolle M, Ashour J, Martínez-Sobrido L, Ashok M, et al. Inhibition of alpha/beta interferon signaling by the NS4B protein of flaviviruses. J Virol 2005;79:8004–8013.

49. Daly C, Reich NC. Double-stranded RNA activates novel factors that bind to the interferon-stimulated response element. Mol Cell Biol 1993;13:3756–3764.

50. Daly C, Reich NC. Characterization of specific DNA-binding factors activated by double-stranded RNA as positive regulators of interferon alpha/beta-stimulated genes. J Biol Chem 1995;270:23739–23746.

51. Weaver BK, Kumar KP, Reich NC. Interferon regulatory factor 3 and CREB-binding protein/p300 are subunits of double-stranded RNA-activated transcription factor DRAF1. Mol Cell Biol 1998;18:1359–1368.

52. Guimarães ES, Gomes MTR, Campos PC, Mansur DS, Dos Santos AA, et al. Cyclic Dinucleotides Trigger STING-Dependent Unfolded Protein Response That Favors Bacterial Replication. J Immunol 2019;202:2671–2681.

53. Antunes KH, Fachi JL, de Paula R, da Silva EF, Pral LP, et al. Microbiota-derived acetate protects against respiratory syncytial virus infection through a GPR43-type 1 interferon response. Nat Commun 2019;10:3273.

54. Ashour J, Laurent-Rolle M, Shi P-Y, García-Sastre A. NS5 of dengue virus mediates STAT2 binding and degradation. J Virol 2009;83:5408–5418.

55. Jones M, Davidson A, Hibbert L, Gruenwald P, Schlaak J, et al. Dengue virus inhibits alpha interferon signaling by reducing STAT2 expression. J Virol 2005;79:5414– 5420.

56. Rand U, Rinas M, Schwerk J, Nöhren G, Linnes M, et al. Multi-layered stochasticity and paracrine signal propagation shape the type-I interferon response. Mol Syst Biol 2012;8:584.

57. Schneider WM, Chevillotte MD, Rice CM. Interferon-stimulated genes: a complex web of host defenses. Annu Rev Immunol 2014;32:513–545.

58. Loeb KR, Haas AL. The interferon-inducible 15-kDa ubiquitin homolog conjugates to intracellular proteins. J Biol Chem 1992;267:7806–7813.

59. Durfee LA, Lyon N, Seo K, Huibregtse JM. The ISG15 conjugation system broadly targets newly synthesized proteins: implications for the antiviral function of ISG15. Mol Cell 2010;38:722–732.

60. Shi H-X, Yang K, Liu X, Liu X-Y, Wei B, et al. Positive regulation of interferon regulatory factor 3 activation by Herc5 via ISG15 modification. Mol Cell Biol 2010;30:2424–2436.

61. Kim M-J, Hwang S-Y, Imaizumi T, Yoo J-Y. Negative feedback regulation of RIG-I-mediated antiviral signaling by interferon-induced ISG15 conjugation. J Virol 2008;82:1474–1483.

62. Okumura F, Okumura AJ, Uematsu K, Hatakeyama S, Zhang D-E, et al. Activation of double-stranded RNA-activated protein kinase (PKR) by interferon-stimulated gene 15 (ISG15) modification down-regulates protein translation. J Biol Chem 2013;288:2839– 2847.

63. D’Cunha J, Knight E Jr, Haas AL, Truitt RL, Borden EC. Immunoregulatory properties of ISG15, an interferon-induced cytokine. Proc Natl Acad Sci U S A 1996;93:211–215.

64. Werneke SW, Schilte C, Rohatgi A, Monte KJ, Michault A, et al. ISG15 is critical in the control of Chikungunya virus infection independent of UbE1L mediated conjugation. PLoS Pathog 2011;7:e1002322.

65. Vuillier F, Li Z, Commere P-H, Dynesen LT, Pellegrini S. USP18 and ISG15 coordinately impact on SKP2 and cell cycle progression. Sci Rep 2019;9:4066.

66. Dabo S, Maillard P, Collados Rodriguez M, Hansen MD, Mazouz S, et al. Inhibition of the inflammatory response to stress by targeting interaction between PKR and its cellular activator PACT. Sci Rep 2017;7:16129.

67. Zanluca C, Mazzarotto GACA, Bordignon J, Duarte Dos Santos CN. Development, characterization and application of monoclonal antibodies against Brazilian Dengue virus isolates. PLoS One 2014;9:e110620.

68. Silveira GF, Wowk PF, Cataneo AHD, Dos Santos PF, Delgobo M, et al. Human T Lymphocytes Are Permissive for Dengue Virus Replication. J Virol;92. Epub ahead of print 15 May 2018. DOI: 10.1128/JVI.02181-17.

69. Mutso M, Saul S, Rausalu K, Susova O, Žusinaite E, et al. Reverse genetic system, genetically stable reporter viruses and packaged subgenomic replicon based on a Brazilian Zika virus isolate. J Gen Virol 2017;98:2712–2724.

70. Osei Kuffour E, Schott K, Jaguva Vasudevan AA, Holler J, Schulz WA, et al. USP18 (UBP43) Abrogates p21-Mediated Inhibition of HIV-1. J Virol;92. Epub ahead of print 15 October 2018. DOI: 10.1128/JVI.00592-18.

