## Supplementary figures and images for "ISG15/USP18/STAT2 is a molecular hub regulating autocrine IFN I-mediated control of Dengue and Zika virus replication"

### Supplemental Figure 1

#### Figure S1-Espada *et al.*

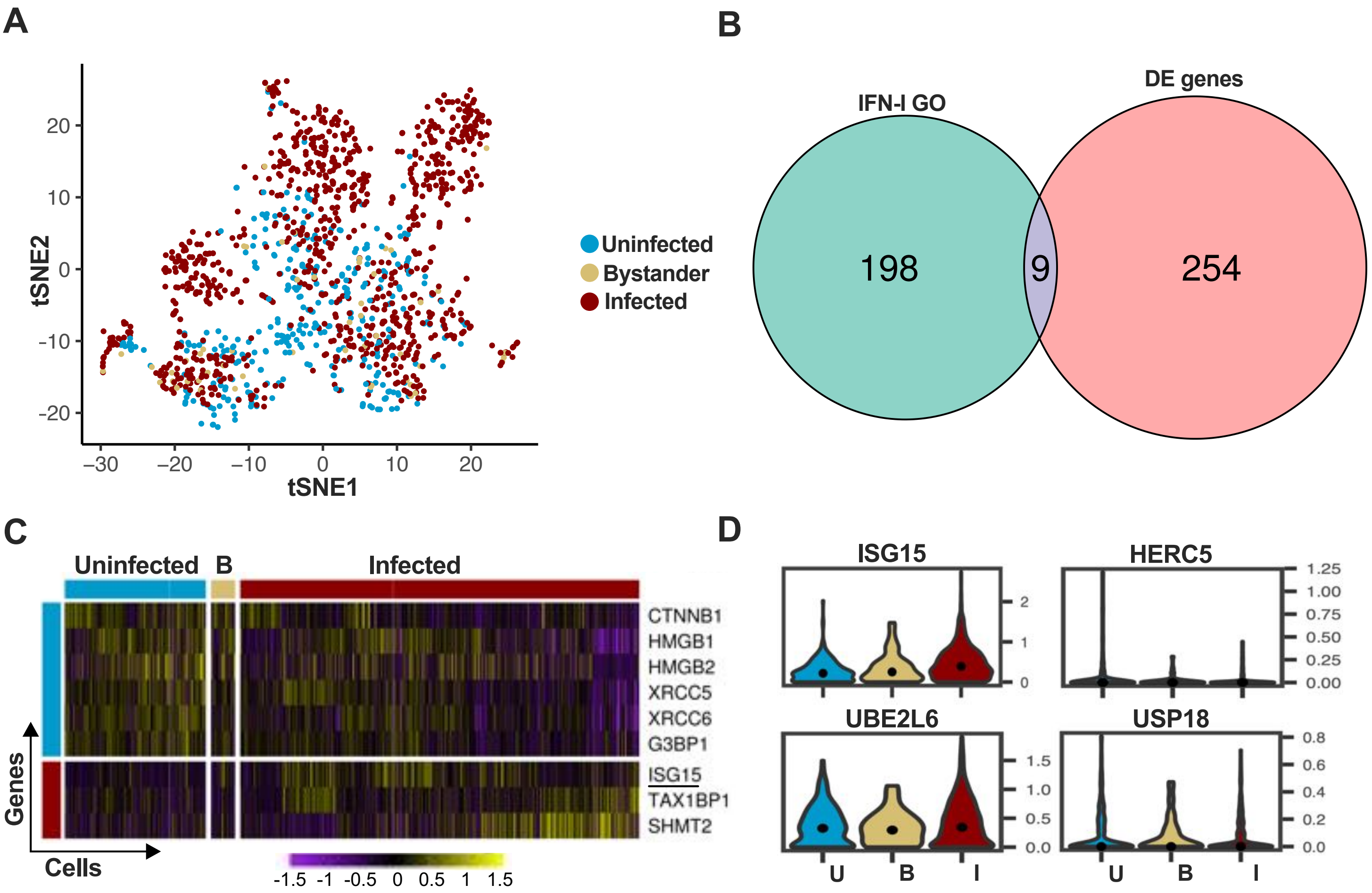

### Supplemental Figure 2

**Figure S2-Espada *et al.***

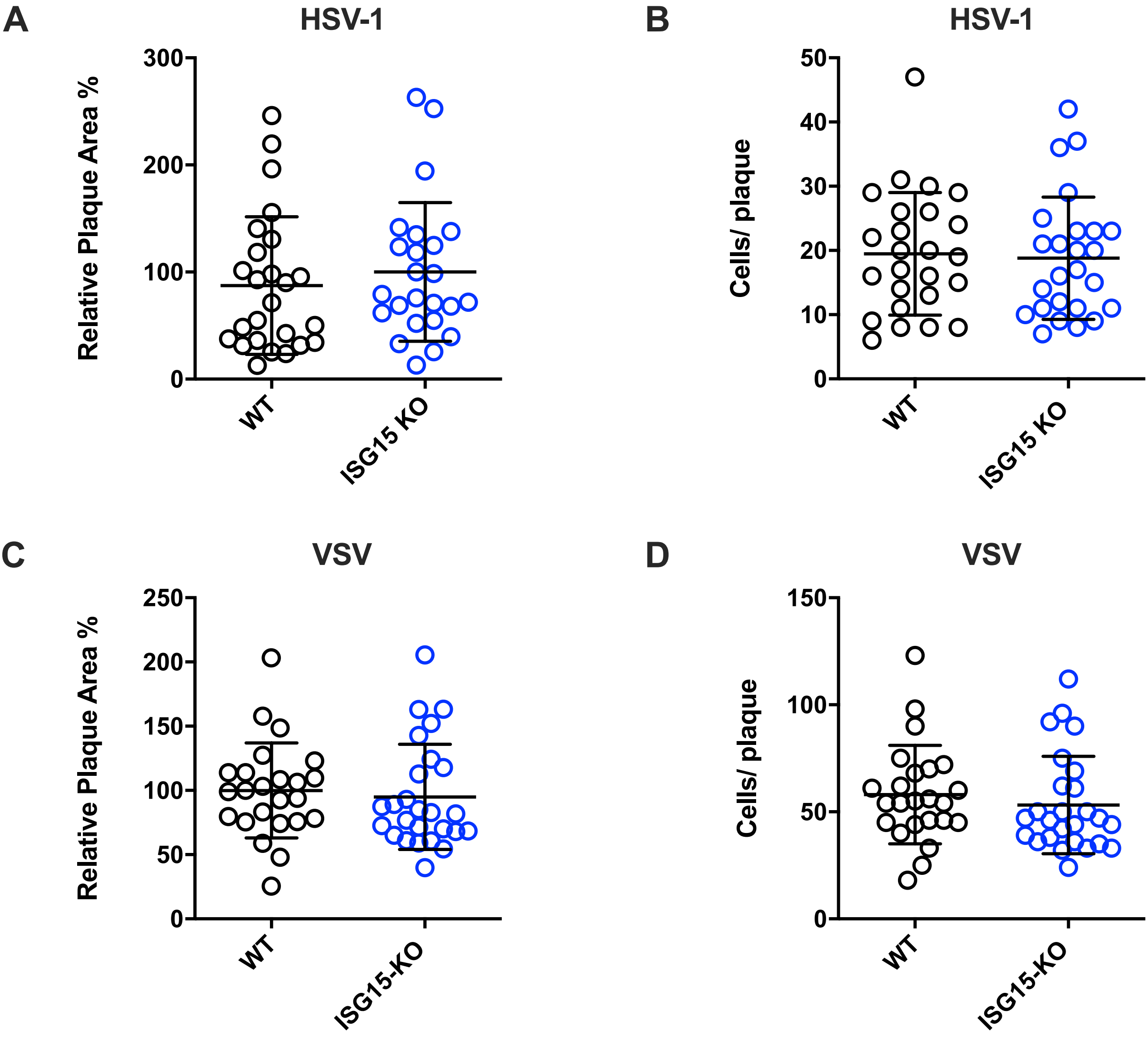

### Supplemental Figure 3

#### Figure S3-Espada *et al.*

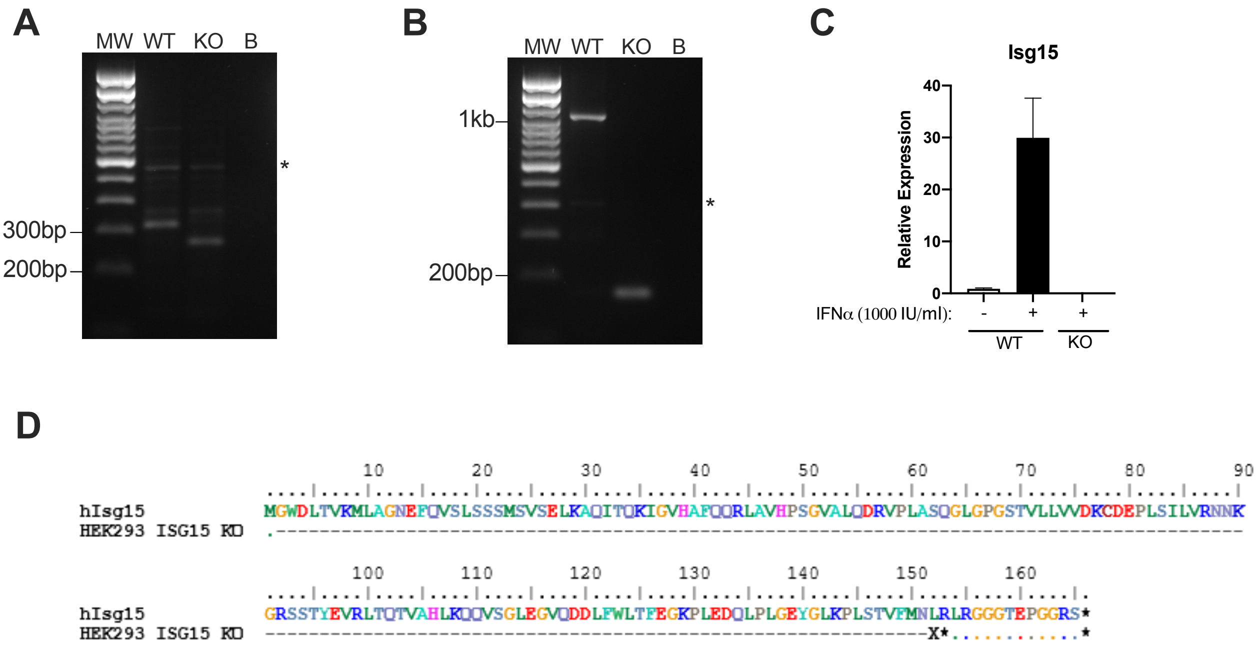

### Supplemental Figure 4

**Figure S4-Espada *et al.***

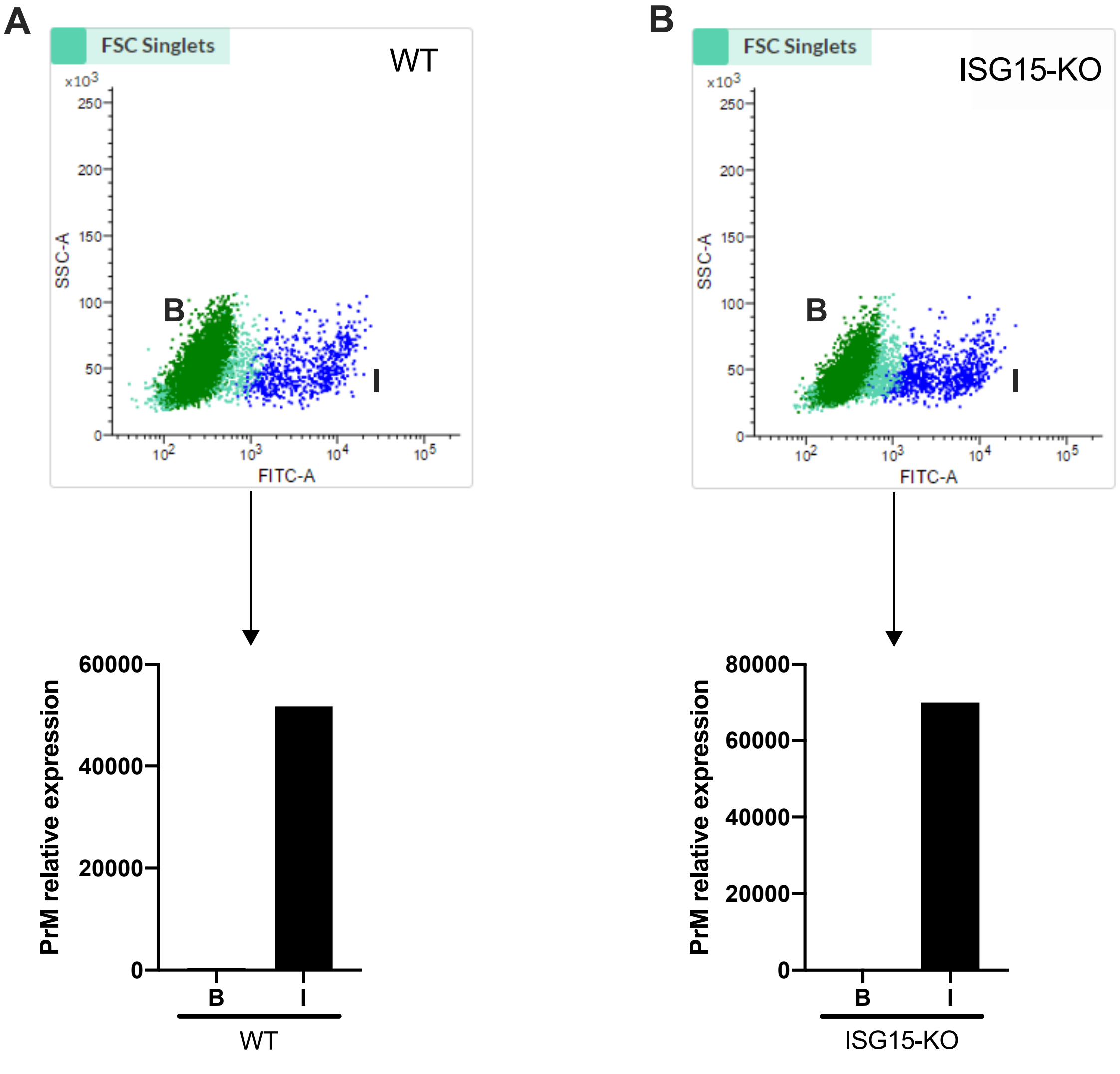
