## Supplemental Table for "ISG15/USP18/STAT2 is a molecular hub regulating autocrine IFN I-mediated control of Dengue and Zika virus replication"

| Supplementary Table 1: sgRNA and primers sequences used in this study | |
| --- | --- |
| Target name | Sequence (5’-) |
| Herc5-gRNA 1 | CGCTCCATCGCCGCTTTGCG |
| Herc5-gRNA 2 | GGCGCGTCACCTCCACCCTC |
| Herc5-PCR genotyping-Fw | CTCTGGGCCTGGGACCC |
| Herc5-PCR genotyping-Rv | CACACACCAGCCCCACTC |
| Ifnar1-gRNA 1 | GCAGCCGCAGGTGAGAGGCG |
| Ifnar1-gRNA 2 | CTGCGGCGGCTCCCAGATGA |
| Ifnar1-PCR genotyping-Fw | CGAACATGTAACTGGTGGGA |
| Ifnar1-PCR genotyping-Rv | CCACTTTCTCCTGGTTGATTTG |
| Isg15-gRNA1 | GGCTGTGGGCTGTGGGCTGT |
| Isg15-gRNA 2 | GGTAAGGCAGATGTCACAGG |
| Isg15-gRNA3 | TGGAGCTGCTCAGGGACACC |
| Isg15KO-PCR genotyping-Fw | GCTGAGAGGCAGCGAACTCAT |
| Isg15KO-PCR genotyping-Rv | GCAGGATCAAGGGCCGGA |
| GFP-Fw | ACCAGCAGAACACCCCCATCG |
| GFP-Rv | GCGGTCACGAACTCCAGCAGG |
| h18S-Fw | TAGAGGGACAAGTGGCGTTC |
| h18S-Rv | CGCTGAGCCAGTCAGTGT |
| DV-PrM-Fw | TTGTCCTAATGATGCTGGTCG |
| DV-PrM-Rv | TCCACCTGAGACTCCTTCCA |
| hIsg15-Fw | TCCTGGTGAGGAATAACAAGGG |
| hIsg15-Rv | GACGACCTGTTCTGGCTGAC |
| hIfit1-Fw | ACGGTATGCTTGGAACGATTG |
| hIfit1-Rv | TGGTGAGGGCTTTCTTTTTCC |
| hUsp18-Fw | CCTGAGGCAAATCTGTCAGTC |
| hUsp18-Rv | CGAACACCTGAATCAAGGAGTTA |
| hIfnB1-Fw | CGAACACCTGAATCAAGGAGTTA |
| hIfnB1-Rv | AGGAGATCTTCAGTTTCGGAGG |

**Table S1- Espada *et al.***
